# Non-Canonical Heme Oxygenase-1 Function in Hematopoietic Stem Cell Homeostasis and Aging

**DOI:** 10.64898/2026.01.26.701757

**Authors:** Monika Zukowska, Izabella Skulimowska, Mateusz Sypniewski, Anna Kusienicka, Mateusz Tomczyk, Sylwester Mosiolek, Jan Morys, Ewa Werner, Karolina Hajduk, Dounia Djeghloul, Michele Goodhardt, Jozef Dulak, Alicja Jozkowicz, Krzysztof Szade

## Abstract

Heme oxygenase-1 (HO-1, encoded by *Hmox1*) is a cytoprotective enzyme with well-established roles in defending against oxidative stress. Global *Hmox1* deficiency in mice accelerates hematopoietic stem cell (HSC) exhaustion and aging, effects previously attributed primarily to loss of HO-1 activity within the bone marrow (BM) niche. However, the cell-intrinsic contribution of HO-1 to HSC regulation has remained unclear.

Here, we show that global *Hmox1* deficiency results in accumulation of an expanded but largely quiescent HSC pool characterized by compromised genome maintenance, altered apoptotic signaling, and defective cell-cycle checkpoint control. We further demonstrate that HO-1 protein is expressed in HSCs and exhibits a predominantly nuclear, non-canonical localization. Using *Hoxb5*-CreERT2-mediated conditional deletion of *Hmox1* in HSCs, we uncover an intrinsic requirement for HO-1 in controlling early hematopoietic differentiation. HSC-specific loss of HO-1 skews stem cell output toward short-term progenitors and increases colony-forming capacity.

Transcriptomic profiling of *Hmox1*^fl/fl^;*Hoxb5*-CreERT2 HSCs revealed broad dysregulation of pathways involved in translation and RNA metabolism, together with aberrant expression of key transcription factors controlling hematopoietic differentiation.

Collectively, these findings identify a non-canonical, cell-intrinsic role for HO-1 in regulating HSC homeostasis, differentiation, and aging.

## Introduction

Heme oxygenase-1 (HO-1) is a critical stress-responsive enzyme that plays a central role in cellular defense mechanisms [1–5]. Traditionally, HO-1 is recognized for its canonical function as the rate-limiting enzyme in heme degradation, catalyzing the oxidative breakdown of heme into carbon monoxide (CO), free iron (Fe2^+^), and biliverdin [6–11]. This enzymatic activity primarily occurs at the smooth endoplasmic reticulum membrane, where HO-1 is typically localized [12]. Through this process, HO-1 contributes to the maintenance of redox homeostasis, cytoprotection, and modulation of inflammatory responses [5,13–16].

Recent evidence, however, suggests that HO-1 exhibits additional roles beyond its classical enzymatic function [17–19]. Under certain stress conditions, HO-1 can translocate to alternative subcellular compartments [20–25], including the nucleus, where it assumes non-canonical functions independent of heme degradation [26–30].

In multiple cell types, a truncated form of HO-1 lacking the C-terminal transmembrane region accumulates in the nucleus. Here, it decreases classical heme-degrading activity but modulates gene regulatory networks [29–32]. Notably, nuclear HO-1 physically interacts with the transcription factor Nrf2, stabilizing it by preventing proteasomal degradation [31]. This stabilization enhances Nrf2 nuclear accumulation and activity, thereby promoting the transcription of detoxification and antioxidant genes in response to oxidative stress [33].

In parallel, nuclear HO-1 has been shown to influence other stress-responsive transcription factors, such as activator protein-1 (AP-1), independent of its enzymatic activity, suggesting broader regulatory effects on gene expression linked to cellular stress responses [27]. Nuclear HO-1 localization has been observed under hypoxia, heme stimulation, and other stress conditions, and has been implicated in modulating transcription factor activity and potentially contributing to disease progression, including cancer [27,34,35]. It has also been proposed that nuclear HO-1 may directly interact with proteins involved in translation, splicing, and DNA damage recognition [36]. We recently demonstrated a modulatory effect of nuclear HO-1 on PARP1 activity and the p53 pathway [37]. Collectively, these findings suggest that HO-1 exerts pleiotropic effects through nuclear interactions, orchestrating transcriptional programs relevant to oxidative stress, inflammation, and cellular adaptation. Importantly, they underscore the need to consider both canonical (enzymatic) and non-canonical (nuclear regulatory) functions of HO-1 in diverse biological systems [38].

Despite its recognized role in the microenvironment, the intrinsic function of HO-1 within hematopoietic stem cells (HSCs) remains incompletely understood. It is known that HO-1 protects hematopoietic stem cells and progenitors from acute stress conditions and regulates myeloid differentiation [39–42]. We have previously shown that HO-1 expression in the bone marrow niche declines with age and that its deficiency accelerates long-term hematopoietic stem cell (LT-HSC) exhaustion [43]. LT-HSCs from young HO-1-deficient mice already display features of premature aging [43]. These observations raised an important question: whether HO-1 deficiency exerts a direct effect on HSC-intrinsic programs or if the observed defects arise exclusively from alterations within the niche.

Here, we demonstrate that the nuclear form of HO-1 is the predominant form in HSCs under steady-state conditions. We show that nuclear HO-1 directly regulates intrinsic HSC functions, including the control of cell cycle and replication stress, and the preservation of genomic integrity. Using a *Hoxb5*-CreER conditional knockout model to delete HO-1 specifically in LT-HSCs, we reveal that nuclear HO-1 deficiency disrupts HSC quiescence and differentiation during aging. These findings establish nuclear HO-1 as a critical intrinsic regulator of HSC function and suggest that targeting nuclear HO-1 pathways may represent a novel strategy to preserve stem cell fitness and counteract age-related hematopoietic decline.

## Results

### HO-1 deficiency expands a pool of quiescent and functionally exhausted HSCs

Global *Hmox1* deficiency accelerates the functional decline of HSCs and promotes premature hematopoietic aging [43]. Global HO-1 knockout (KO) mice already exhibit an expanded HSC compartment with a higher proportion of cells in active phases of the cell cycle. However, it remained unclear whether the entire expanded HSC pool actively proliferates and sustains the accelerated hematopoiesis observed in HO-1 KO mice.

To assess the proliferative dynamics of the HSC pool over time, we performed an *in vivo* BrdU incorporation assay, consisting of an initial intraperitoneal BrdU injection followed by continuous BrdU administration in drinking water to HO-1 KO and wild-type (WT) mice (Fig. 1A). Long-term, continuous BrdU labeling revealed markedly lower overall incorporation in HO-1 KO HSCs compared with WT controls (Fig. 1B). In WT mice, the fraction of BrdU^+^ HSCs progressively increased, reaching saturation at 60–70%, indicating that most HSCs underwent at least one division during the labeling period and that there is a constant influx of dividing cells into the pool. In contrast, in HO-1 KO mice, the proportion of BrdU^+^ HSCs plateaued at approximately 20%, indicating that only a small subset of HSCs actively divides, while the majority remain dormant over extended periods. A similar pattern was observed in the broader LKS (Lineage^-^c-Kit^+^Sca-1^+^) progenitor fraction (Fig. 1C), suggesting that a large portion of hematopoiesis in HO-1 KO mice is relatively inactive and rarely incorporates BrdU. These data imply that blood cells production in HO-1 KO mice relies on a minor fraction of hematopoietic cells compared with WT mice.

**Figure 1.**
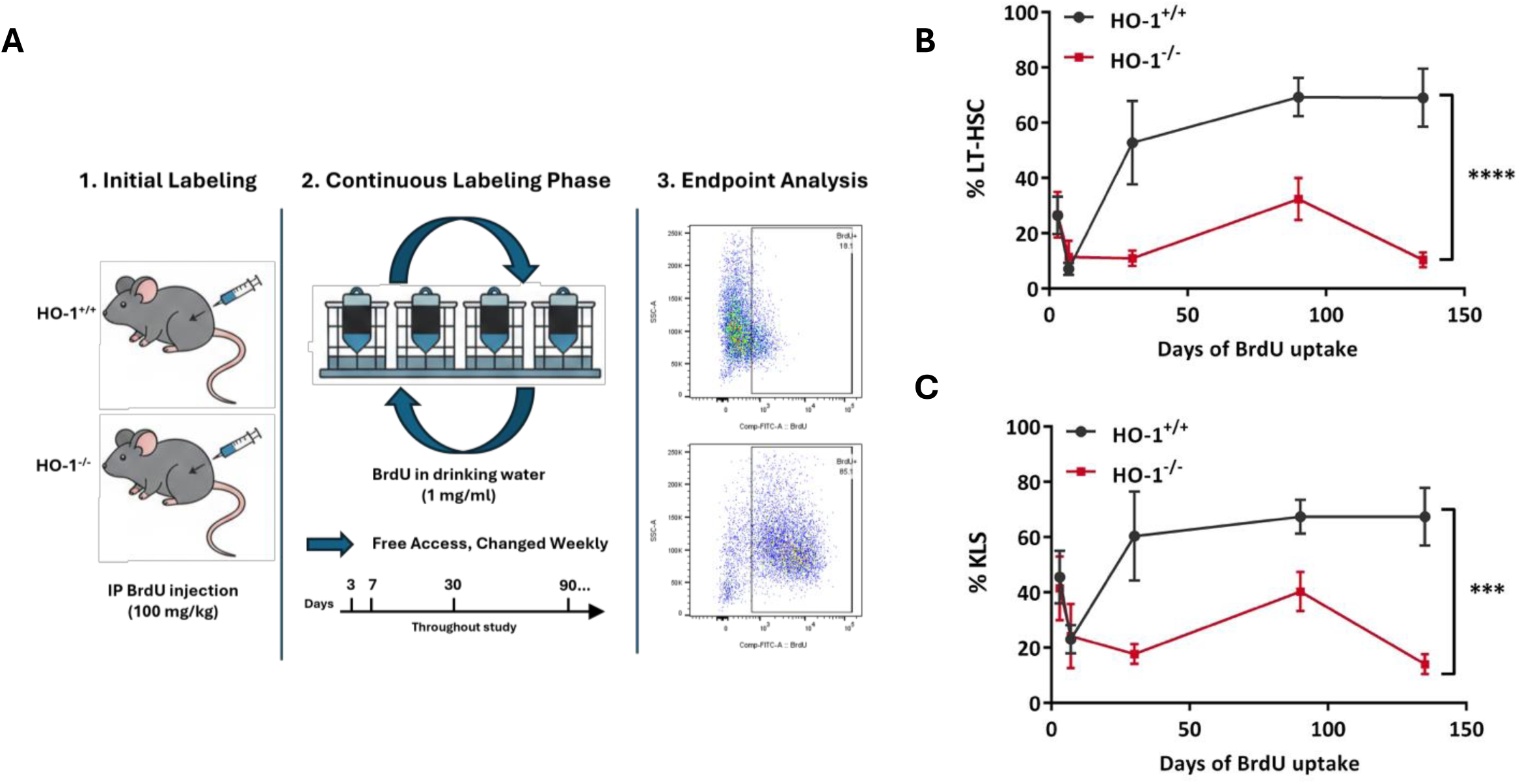
HO-1 deficiency expands a pool of quiescent HSCs with limited proliferative activity. (A) Experimental scheme of long-term *in vivo* BrdU incorporation assay. Wild-type (WT) and HO-1 knockout (HO-1 KO) mice received an initial intraperitoneal BrdU injection followed by continuous BrdU administration in drinking water. The percentage of BrdU^+^ HSCs was quantified over time by flow cytometry. (B) Percentage of BrdU^+^ cells within the LT-HSC pool during long-term labeling in WT and HO-1 KO mice. (C) Percentage of BrdU^+^ cells within the KLS (Lin^−^Sca-1^+^c-Kit^+^) progenitor fraction during long-term labeling in WT and HO-1 KO mice. Data show mean ± SEM, n = 5-6 mice/group. Statistical significance was assessed using 2-way ANOVA.

Together, these results indicate that HO-1 deficiency leads to an expanded but functionally heterogeneous HSC compartment, composed predominantly of long-term quiescent cells, with only a small subset actively contributing to hematopoiesis. This functional heterogeneity may underlie the premature exhaustion of the HSC pool in HO-1-deficient mice.

### HO-1–deficient HSCs accumulate DNA damage

Previously, we demonstrated that young HO-1^-/-^ HSCs exhibited significantly elevated DNA damage, as reflected by increased tail DNA content and olive tail moment compared with age-matched WT controls in alkaline comet assay [43].

Given the heterogenous proliferative activity within the aged old HO-1^-/-^ HSC pool, we performed a similar analysis in old mice (Fig. 2A). Interestingly, aged HO-1^-/-^ HSCs showed lower levels of DNA breaks than age-matched WT controls (Fig. 2A). This observation is consistent with the predominance of long-term quiescent cells in the HO-1^-/-^ HSC compartment in aged mice, which may limit the accumulation of replication-associated DNA damage.

**Figure 2.**
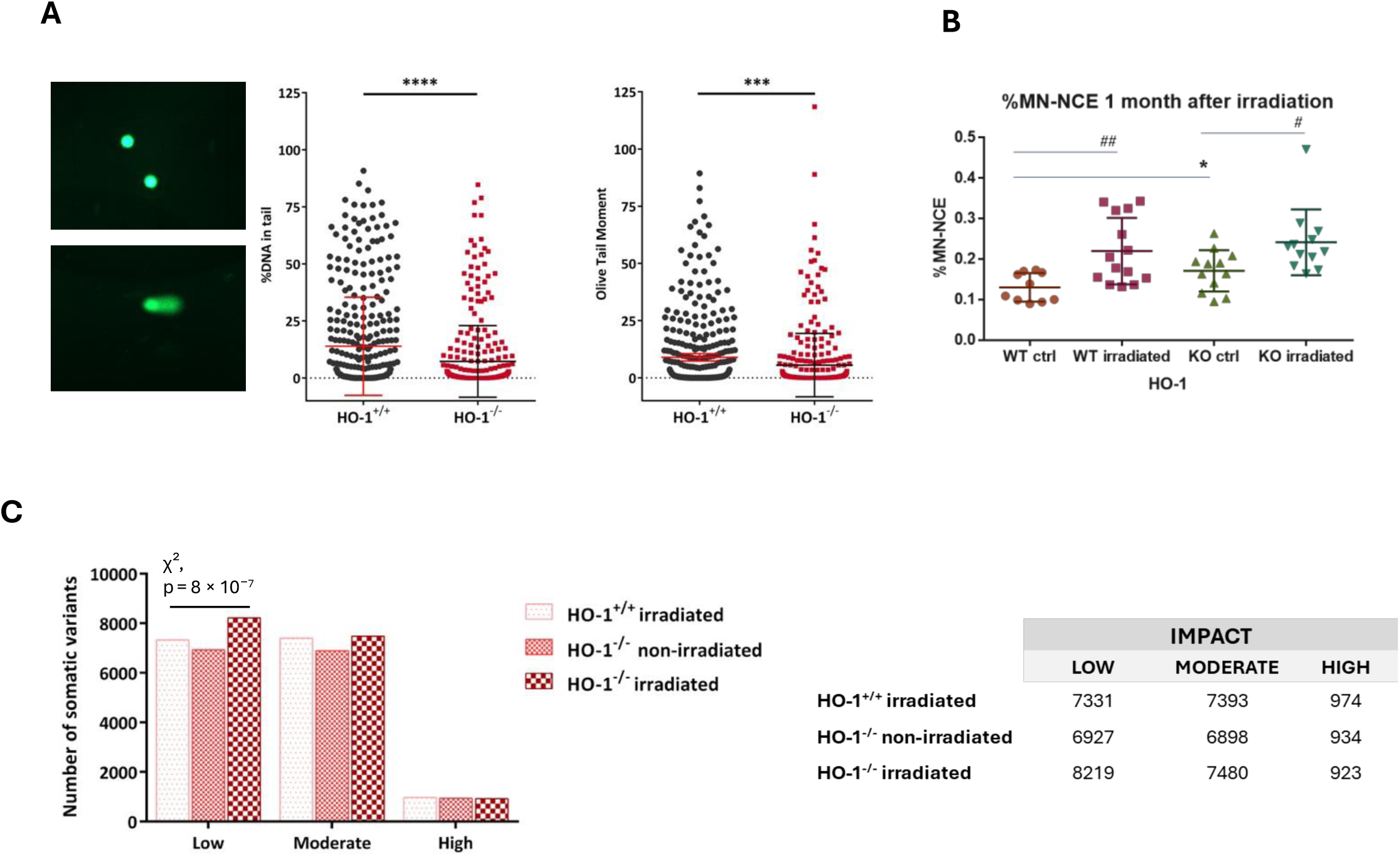
HO-1–deficient HSCs accumulate DNA damage and show altered mutational outcomes after genotoxic stress. (A) Quantification of DNA damage in aged WT and HO-1 KO HSCs. Data are shown as mean ± SD. n = 334-368 cells from pooled 3-4/mice per group. (B) Quantification of micronuclei in peripheral blood normochromatic erythrocytes (NCE) indicates increased chromosomal damage in HO-1 KO HSCs in steady state conditions and after irradiation during erythropoiesis. Points represent individual mice. (C) Distribution of mutation-impact categories (low, moderate, high impact) identified by whole-bone-marrow exome sequencing six months after 2 Gy irradiation in WT and HO-1 KO mice. Statistical significance assessed by chi-square test of independence.

To assess whether HO-1 deficiency alters the cellular response to genotoxic stress and the long-term maintenance of genome integrity in hematopoietic cells, we exposed young 3-month-old WT and HO-1^−/−^ mice to low-dose irradiation (2 Gy). One month after irradiation, we evaluated micronuclei in peripheral blood normochromatic erythrocytes (NCE), providing an integrated readout of chromosomal damage accumulated during erythropoiesis (Fig. 2B). Under control conditions, the fraction of cells harboring micronuclei was low but detectable. In both genotypes, this fraction increased significantly following irradiation. Notably, under control conditions, micronuclei were more frequently detected in HO-1^−/−^ mice. This finding may indicate an increased susceptibility to chromosomal instability, resulting from replication stress or segregation-associated defects in HO-1 deficient cells, which are commonly reflected by micronucleus formation. After irradiation, micronucleus frequencies were comparable between genotypes, suggesting that under conditions of acute genotoxic stress, HO-1 deficiency does not further exacerbate the extent of chromosomal damage detectable by this assay.

To assess the long-term consequences of genotoxic stress across the hematopoietic landscape, mice were allowed a 6-month recovery period post-irradiation, after which we performed whole-bone marrow exome sequencing, using non-irradiated WT littermates as a germline reference. This approach enabled the identification of persistent mutational events, which were classified as low-, moderate-, or high-impact variants (Fig. 2C) [44].

Most detected variants in both genotypes were classified as low- or moderate-impact mutations, such as missense substitutions (Supplementary Table 1). Importantly, the distribution of mutation-impact categories differed significantly between WT and HO-1^-/-^ mice following irradiation (p = 2 × 10^−^⁶, chi-square test of independence, Fig. 2C). This shift was driven primarily by an increase in low-impact mutations in HO-1^-/-^ animals (p = 8.05 × 10^−^⁷), whereas moderate- and high-impact variants were largely comparable between genotypes (Fig. 2C). This pattern aligns with the expectation that irradiation-induced moderate or high-impact mutations often lead to cell elimination, leaving predominantly low-impact mutations detectable in surviving long-lived cells [45,46].

Collectively, these findings demonstrate that HO-1 deficiency compromises genome maintenance and increases susceptibility to mutagenic stress in young HSCs. However, consistent with our BrdU-based proliferation analysis, the HSC compartment in aged HO-1^-/-^ mice is largely composed of non-cycling cells, which may paradoxically protect it from proliferation-associated DNA damage despite an impaired repair environment.

### HO-1 deficiency disrupts DNA damage checkpoint control

Given the elevated DNA damage already present in young HO-1 KO HSCs, we next examined signaling pathways involved in the DNA damage response and cell cycle regulation.

Annexin V staining revealed that, despite their increased DNA damage, young HO-1 KO HSCs did not show elevated apoptosis (Fig. 3A). In contrast, WT HSCs exhibited a higher proportion of early apoptotic cells, suggesting that HO-1–deficient HSCs fail to properly activate apoptosis in response to DNA damage (Fig. 3A). The same was true for the broader KLS fraction, with more viable cells and fewer early apoptotic cells in HO-1^-/-^ animals (Fig. 3B).

**Figure 3.**
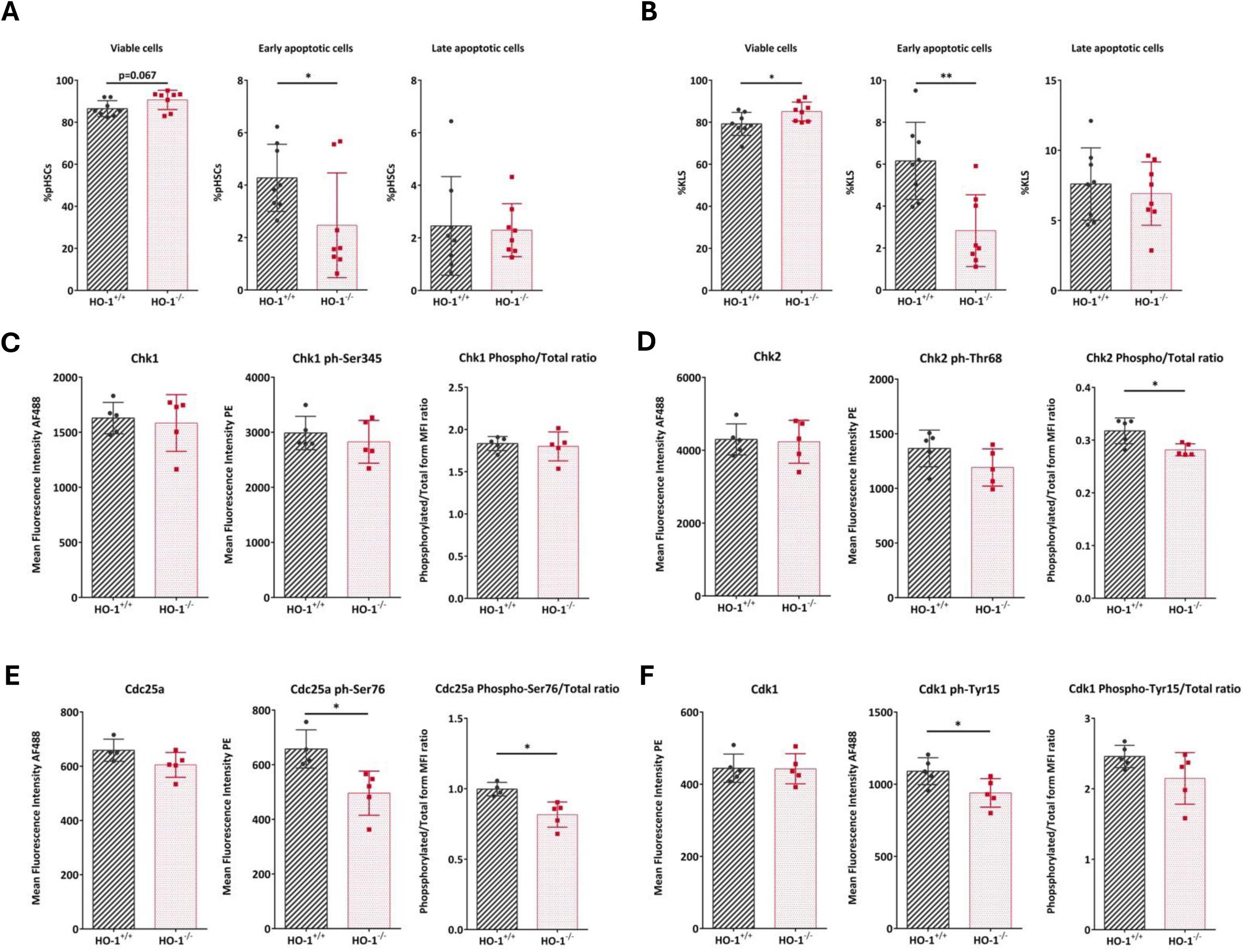
HO-1 deficiency impairs DNA damage-induced checkpoint activation and apoptosis in HSCs. (A) Annexin V assay results comparing the percentage of viable and apoptotic cells between HO-1 WT and HO-1 KO HSCs, and (B) HO-1 WT and HO-1 KO KLS cells. (C) Flow cytometry analysis of Chk1 expression and its phosphorylated form in HSCs, together with their ratio in both genotypes. (D) Flow cytometry analysis of Chk2 expression and its phosphorylated form in HSCs, together with their ratio in both genotypes. (E) Flow cytometry analysis of Cdc25a expression and its phosphorylated form in HSCs, together with their ratio in both genotypes. (F) Flow cytometry analysis of Cdk1 expression and its phosphorylated form in HSCs, together with their ratio in both genotypes. Points represent individual mice. Data are shown as mean ± SD.

Therefore, we next examined the status of checkpoint kinases Chk1 and Chk2, key mediators of genome-integrity checkpoints [47]. Although total and phosphorylated Chk1 and Chk2 levels were broadly similar between groups (Fig. 3C, D), the ratio of phosphorylated to total Chk2 was moderately but significantly increased in HO-1 KO HSCs (Fig. 3D), indicating impaired Chk2 activation and a potential failure to induce cell cycle arrest upon DNA damage. Consistent with this, phosphorylation of the Chk2 target Cdc25A at Ser76 – a modification promoting its degradation and enforcing the G1/S checkpoint [48] – was reduced in HO-1 KO HSCs (Fig. 3E).

Downstream of Cdc25A, we evaluated Cdk1, a cyclin-dependent kinase whose activity is tightly controlled by Tyr15 phosphorylation [49,50]. HO-1 KO HSCs exhibited reduced phosphorylation of Cdk1 at Tyr15 compared with WT cells, although total Cdk1 and phosphorylated/total ratios did not reach statistical significance (Fig. 3F).

Next, we assessed Cdc45, a replication helicase activated by Cdk1 and Cdk2 and required for origin firing [51,52]. Although average Cdc45 levels per mouse did not differ between genotypes, flow cytometry analysis revealed a greater proportion of Cdc45-high HSCs in HO-1 KO mice, suggesting aberrant activation of replication initiation in a subset of cells (Fig. 4A).

**Figure 4.**
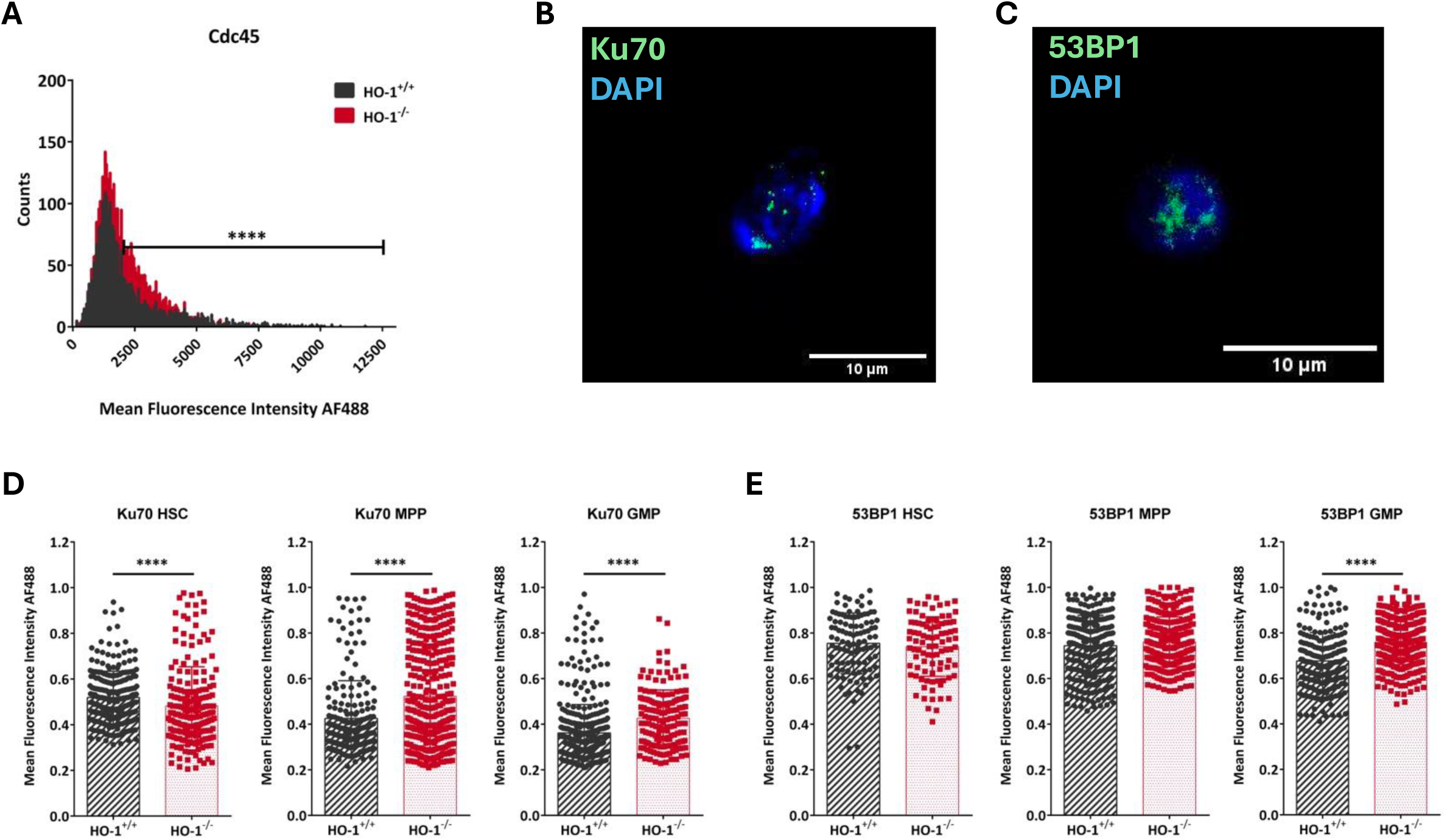
HO-1 deficiency disrupts early DNA damage sensing selectively in HSCs. (A) Flow cytometry analysis of Cdc45 expression in WT and HO-1 KO HSCs, showing the proportion of Cdc45-high cells. n = 5-6 mice/group. For statistical significance calculations in signals obtained from individual n = 755-1157 cells from pooled mice were taken into account. (B, C) Representative confocal images of Ku70 and 53BP1 nuclear foci in sorted HSCs. (D) Quantification of Ku70 nuclear signal intensity in HSCs, MPPs, and GMPs from WT and HO-1 KO mice. n = 25-50 cells/group from pooled 3 mice per genotype. (E) Quantification of 53BP1 nuclear signal intensity in HSCs, MPPs, and GMPs. Each dot represents an individual cell, cells were pooled from multiple mice. Bars indicate mean ± SEM.

Altogether, these results indicate that HO-1 deficiency disrupts multiple components of the G1/S checkpoint, including Chk2 activation, Cdc25A regulation, Cdk1 phosphorylation, and replication origin licensing.

Finally, to explore a potential mechanistic link between increased DNA damage and defective checkpoint activation in HO-1 KO HSCs, we examined the DNA damage-sensing proteins Ku70 and 53BP1. These factors participate in early recognition of DNA breaks and help recruit or activate downstream checkpoint components, including the Chk2/Cdc25A axis [53,54]. Using confocal microscopy, we quantified both the number and fluorescence intensity of Ku70 and 53BP1 nuclear foci in sorted HSCs, MPPs, and GMPs (Fig. 4B, C).

In HSCs from HO-1-deficient mice Ku70 signal intensity was significantly reduced, indicating impaired recruitment or stabilization of Ku70 at DNA lesions (Fig. 4D). Strikingly, the opposite trend was observed in more differentiated progenitors: Ku70 intensity was increased in MPPs and GMPs from HO-1 KO mice (Fig. 4D). For 53BP1, we detected no changes in HSCs or MPPs; however, GMPs from HO-1 KO mice exhibited significantly higher 53BP1 intensity (Fig. 4E).

These findings point to an HSC-specific defect in early DNA-damage sensing in the absence of HO-1. The reduced Ku70 signal in HO-1 KO HSCs, despite their elevated levels of DNA damage, may contribute to the failure to enforce G1/S checkpoint arrest. In contrast, the increased Ku70 and 53BP1 signals in progenitors suggest that HO-1 may regulate DNA-damage responses differently across the hematopoietic hierarchy.

Altogether, these results support a model in which HO-1 deficiency disrupts DNA-damage checkpoint control specifically in HSCs by weakening the earliest steps linked to double-strand break detection.

### HO-1 shows non-canonical nuclear localization in HSCs

Our previous research found that while HO-1 expression in the bone marrow is the highest in endothelial cells, mesenchymal cells, and macrophages, it is also detectable in hematopoietic stem and progenitor cells [43]. Therefore, we analyzed the expression of HO-1 in early stages of hematopoietic differentiation. HO-1 mRNA expression in stem cells and multipotent progenitors in young (3-month-old) mice increases with differentiation from LT-HSCs (KLS Flt3^-^CD150^+^CD34^-^) to ST-HSCs (KLS Flt3^-^CD34^+^) and multipotent progenitors (MPPs - KLS Flt3^+^CD34^+^). Interestingly, in LT-HSCs, HO-1 level is similar in young and old mice (18-month-old), but in ST-HSCs and MPPs, its expression decreases with age and is significantly lower in aged animals (Fig. 5A).

**Figure 5.**
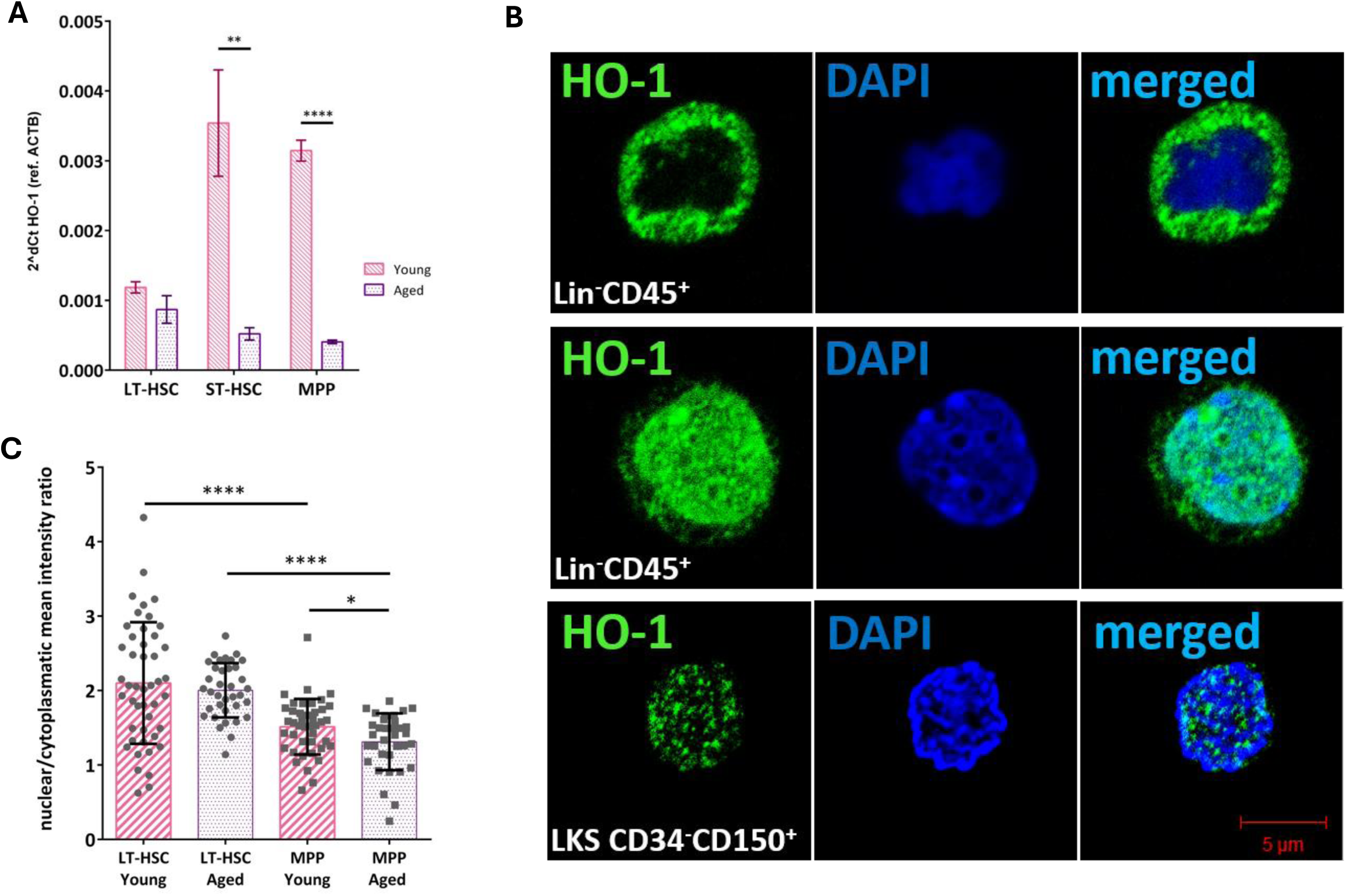
HO-1 exhibits non-canonical nuclear localization in LT-HSCs. (A) HO-1 mRNA expression levels in LT-HSCs, ST-HSCs, and MPPs isolated from young (3-month-old) and aged (18-month-old) mice. (B) Representative confocal images showing cytoplasmic versus nuclear localization of HO-1 protein in Lin^−^CD45^+^ bone marrow cells and LT-HSCs. (C) Quantification of HO-1 subcellular localization in LT-HSCs and MPPs from young and aged mice. Data are shown as mean ± SD. n = 33-45 cells/group

We next sought to determine the intracellular localization of HO-1 protein in hematopoietic cells. Within the Lin^−^CD45^+^ bone marrow compartment, we identified cells with conventional cytoplasmic HO-1 expression, as well as cells in which HO-1 was strictly nuclear (Fig. 5B). Strikingly, LT-HSCs (LKS CD34^-^CD150^+^) displayed predominantly nuclear HO-1 localization, often forming discrete foci-like structures, suggesting a non-canonical function of HO-1 in the nuclear compartment of these cells (Fig. 5B).

We checked whether such non-classical nuclear localization is limited to hematopoietic stem cells or if it can also be observed in hematopoietic progenitors. Therefore, we analyzed the localization of HO-1 protein in LT-HSCs and MPPs in young and aged mice. We found that the nuclear localization of HO-1 is indeed characteristic of both young and old LT-HSCs, whereas in MPPs, HO-1 had more cytoplasmic localization, especially in aged cell donors (Fig. 5C).

### HSC-intrinsic deletion of HO-1 rewires transcriptional programs governing activation and lineage priming

Expression of HO-1 within the hematopoietic niche is essential for maintaining full HSC functionality. Our data demonstrated that global HO-1 deficiency leads to increased DNA damage and impaired cell cycle checkpoint activation. To determine whether HO-1 also plays a cell-intrinsic regulatory role within HSCs, we generated *Hoxb5*-CreERT2;*Hmox1*^fl/fl^ mice, enabling inducible and LT-HSC–specific deletion of *Hmox1*. Young mice were treated with tamoxifen to activate Cre recombinase and subsequently aged for one year under physiological, unperturbed conditions. LT-HSCs were then sorted from *Hoxb5*-CreERT2;*Hmox1*^fl/fl^ mice (HO-1^ΔHSC^) and control *Hoxb5*-CreERT2;*Hmox1*^WT^ littermates (WT), and their transcriptomes were analyzed by RNA sequencing.

Differential expression analysis identified 492 genes significantly upregulated and 219 genes downregulated in HO-1^ΔHSC^ LT-HSCs compared with WT controls (FDR < 0.1, n = 3–5 samples per group, each sample pooled from two mice). These findings demonstrate that loss of HO-1 within HSCs, despite their residence in an otherwise HO-1–competent niche, substantially alters their transcriptional landscape (Fig. 6A, B).

**Figure 6.**
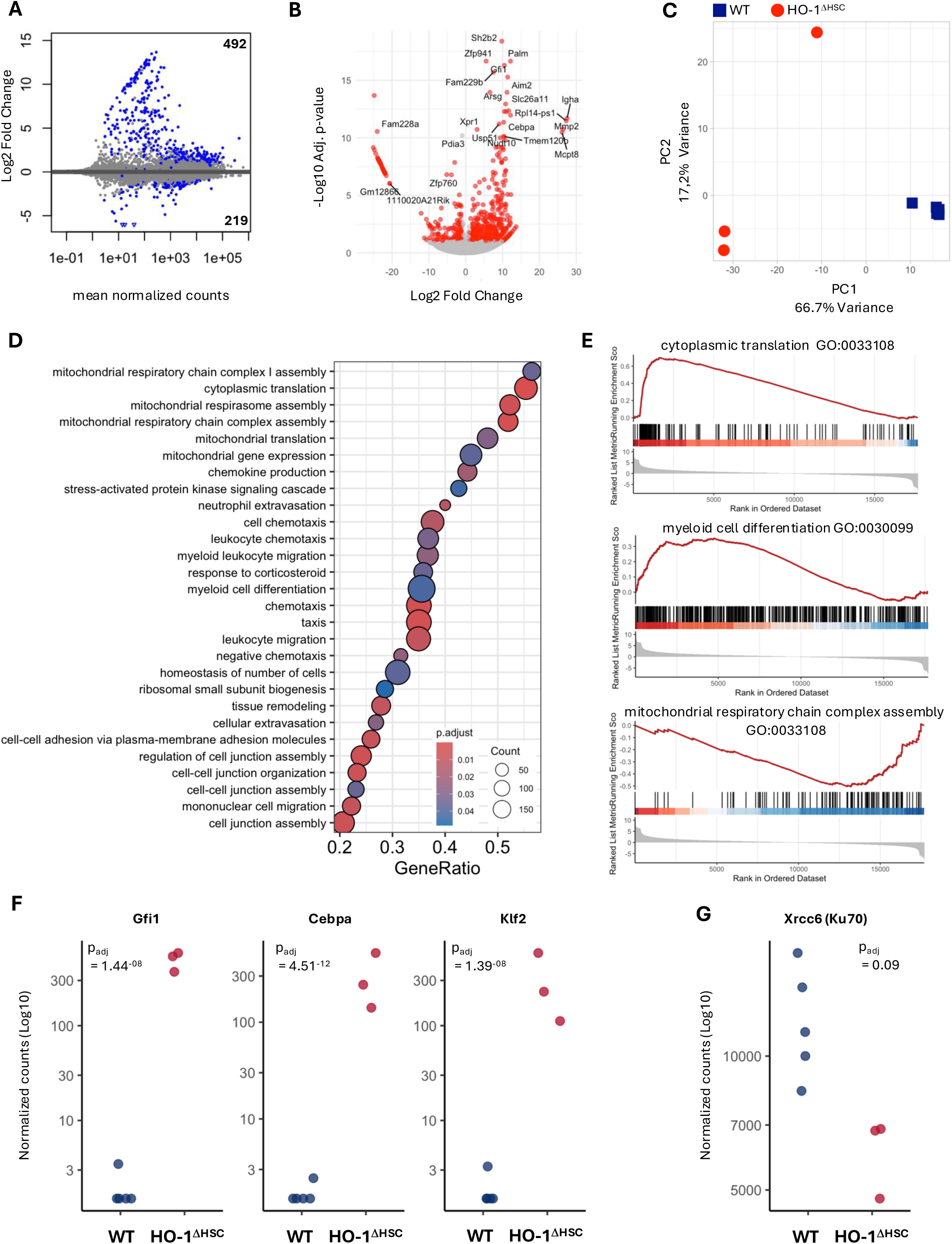
HSC-intrinsic deletion of HO-1 rewires transcriptional programs linked to activation and lineage priming. (A) MA plot showing the number of differentially expressed genes in LT-HSCs isolated from HO-1^ΔHSC^ and control mice. (B) Volcano plot showing differentially expressed genes in LT-HSCs isolated from HO-1^ΔHSC^ and control mice. (C) Principal component analysis (PCA) of WT and HO-1^ΔHSC^ LT-HSC transcriptomes. (D) Overview of significantly enriched Gene Ontology (GO) biological processes identified by gene set enrichment analysis (GSEA). (E) Representative GSEA enrichment plots for cytoplasmic translation, myeloid differentiation, and mitochondrial respiration pathways. (F) Expression levels of selected transcription factors regulating hematopoietic fate decisions (Gfi1, Cebpa, Klf2). (G) Expression of Xrcc6 (Ku70) in WT and HO-1^ΔHSC^ LT-HSCs.

Principal component analysis (PCA) based on differentially expressed genes revealed clear segregation between WT and HO-1^ΔHSC^ samples, with PC1 accounting for 66.7% of the variance and effectively separating genotypes. PC2 explained 17.2% of the variance and primarily reflected heterogeneity within the HO-1^ΔHSC^ group (Fig. 6C).

To identify biological processes associated with these transcriptional changes, we performed gene set enrichment analysis (GSEA). Among significantly enriched Gene Ontology (GO) biological processes (Supplementary Fig. 1), we observed prominent signatures related to cytoplasmic translation, mitochondrial respiration, and leukocyte biology, including differentiation, migration, and adhesion (Fig. 6D). Notably, cytoplasmic translation pathways were strongly and positively enriched in HO-1^ΔHSC^ LT-HSCs (Fig. 6E; Supplementary Fig. 2), consistent with increased cellular activation. In parallel, gene sets associated with myeloid differentiation were also positively enriched, suggesting a potential myeloid bias in HO-1–deficient LT-HSCs (Fig. 6E). In contrast, pathways related to mitochondrial respiration showed significant negative enrichment (Fig. 6E; Supplementary Fig. 2).

Inspection of individual genes contributing to the myeloid differentiation signature revealed aberrant expression of key transcription factors governing hematopoietic fate decisions. The myeloid regulators Gfi1 and Cebpa were largely absent in WT LT-HSCs but consistently expressed in HO-1^ΔHSC^ cells (Fig. 6F). Interestingly, we also observed elevated expression of Klf2 specifically in HO-1^ΔHSC^ LT-HSCs, a transcription factor previously shown to restrain monocyte differentiation and myeloid cell activity (Fig. 6F). Together, these findings suggest that aged HO-1^ΔHSC^ LT-HSCs exhibit features of mixed or aberrant lineage priming rather than a unidirectional differentiation program.

Finally, we assessed whether genes involved in cell cycle regulation and DNA damage responses, previously altered in HSCs from global HO-1–deficient mice (Fig. 3 and Fig. 4), were also affected in HO-1^ΔHSC^ LT-HSCs. While we did not detect changes in the expression of cyclin-dependent kinases, whose activity is largely regulated post-translationally [55], we observed reduced expression of Xrcc6 (encoding Ku70) in HO-1^ΔHSC^ LT-HSCs (Fig. 6G). This reduction mirrors findings in global HO-1–deficient HSCs (Fig. 4D), suggesting that diminished Ku70 expression may represent a direct, cell-intrinsic consequence of HO-1 loss. However, consistent with our GSEA results, we did not observe significant enrichment of broader DNA damage response pathways.

### Intrinsic HO-1 deficiency promotes early LT-HSC commitment without altering lineage output

We next examined how HO-1 deletion within HSCs affects their functional behavior. Flow cytometric analysis revealed that conditional deletion of *Hmox1* in LT-HSCs did not significantly alter the overall frequencies of LT-HSCs or short-term HSCs (ST-HSCs) (Fig. 7A). However, the LT-HSC/ST-HSC ratio was significantly reduced (Fig. 7C), indicating a shift toward early differentiation and commitment of LT-HSCs. Consistently, we observed an increased frequency of ST-HSCs within the LSK progenitor fraction (Fig. 7B), further supporting enhanced early activation and differentiation upon intrinsic HO-1 loss.

**Figure 7.**
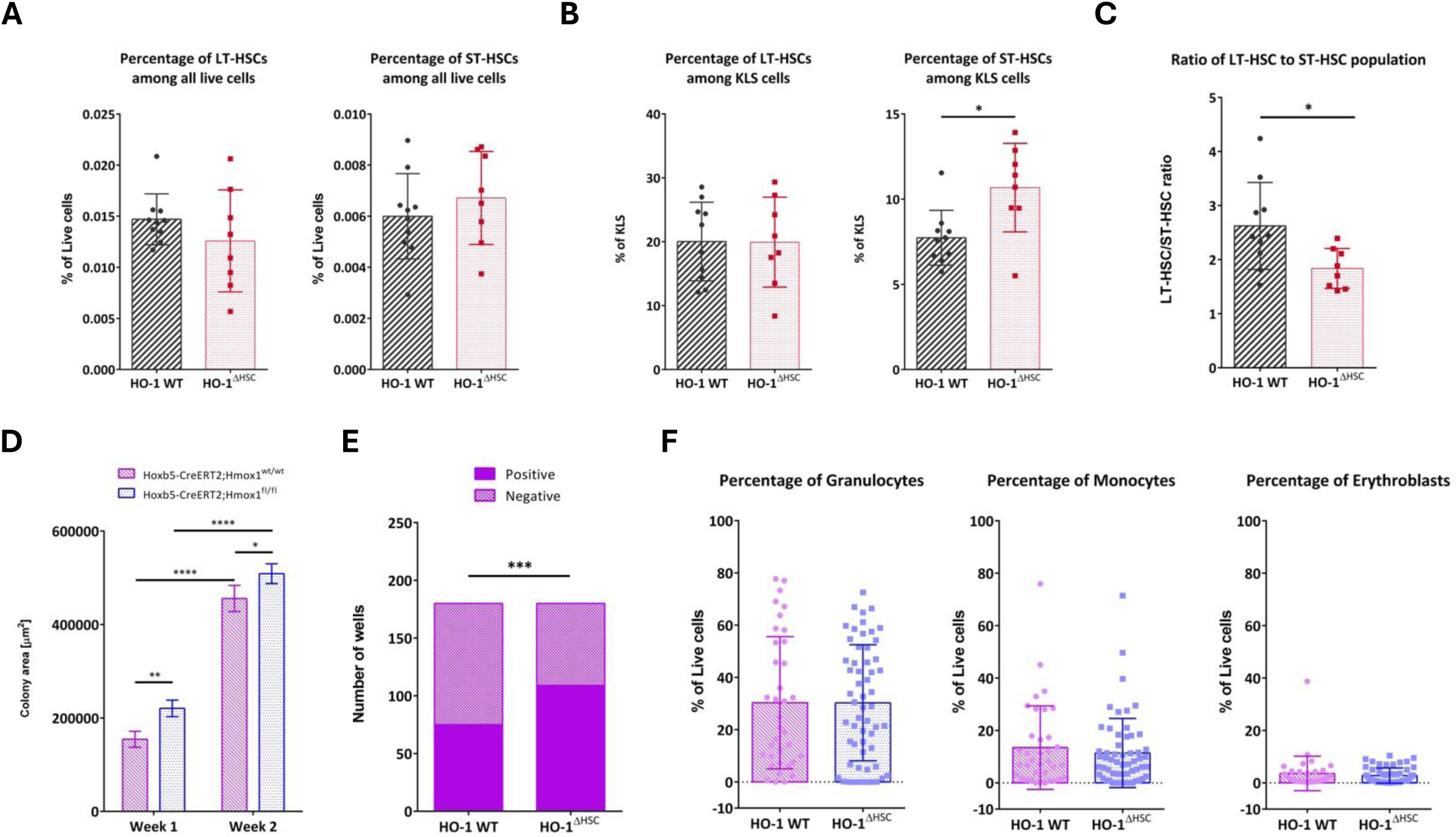
Intrinsic HO-1 deficiency promotes early LT-HSC commitment without altering lineage output. (A) Flow cytometric quantification of LT-HSC and ST-HSC frequencies in WT and HO-1^ΔHSC^ mice. (B) LT-HSC/ST-HSC ratio in WT and HO-1^ΔHSC^ mice. (C) Frequency of ST-HSCs within the LSK progenitor fraction. Data are shown as mean ± SD. (D) Colony growth kinetics from single LT-HSCs isolated from WT and HO-1^ΔHSC^ mice. (E) Frequency of formed colonies from single LT-HSCs. Data are shown as mean ± SEM. (F) Flow cytometric analysis of lineage composition of colonies derived from WT and HO-1^ΔHSC^ LT-HSCs. Data are shown as mean ± SD. Each dot represents an individual clone or mouse, as indicated.

To functionally validate the activation, priming, and differentiation signatures observed at the transcriptomic and phenotypic levels, we performed a single-cell *in vitro* differentiation assay. Individual LT-HSCs were sorted into differentiation-permissive conditions, and clonal growth was monitored over three weeks. At the end of the assay, the cellular composition of individual colonies was analyzed by flow cytometry. This approach enabled high-throughput functional assessment at the clonal level, which is particularly important given that bulk RNA-sequencing signatures may reflect heterogeneous lineage priming across distinct HSC clones.

Consistent with enhanced activation, LT-HSCs from HO-1^ΔHSC^ mice formed colonies more rapidly during the first and second week of culture (Fig. 7D), indicating an accelerated transition from a quiescent stem cell state toward proliferating progenitors. By the end of the assay, HO-1^ΔHSC^ LT-HSCs also exhibited increased colony-forming efficiency compared with WT controls (Fig. 7E).

Despite these differences in activation kinetics and clonogenic output, the lineage composition of colonies was not altered (Fig. 7F). Myelopoiesis remained the predominant differentiation outcome in both genotypes (Fig. 7F), indicating that intrinsic HO-1 loss enhances early differentiation potential without overtly skewing lineage fate decisions under these conditions.

Together, these data demonstrate that HO-1 acts intrinsically within HSCs to restrain early activation and differentiation, thereby contributing to the maintenance of the LT-HSC state.

## Discussion

In this study, we identify a previously unrecognized, non-canonical, and cell-intrinsic role for HO-1 in the regulation of HSC homeostasis, differentiation, and aging. While HO-1 has long been appreciated as a cytoprotective, stress-inducible enzyme acting primarily through heme degradation and redox control [3,4], our findings demonstrate that HO-1 also plays an HSC-intrinsic role. HO-1 localizes predominantly to the nucleus of HSCs and modulates stem cell function.

A key conceptual advance of our work is the demonstration that global HO-1 deficiency does not simply drive generalized hyperproliferation of the HSC compartment. Instead, using long-term BrdU labeling, we reveal that HO-1–deficient mice harbor an expanded but functionally heterogeneous HSC pool, in which only a small fraction of stem and progenitor cells actively cycle and sustain hematopoiesis (Fig. 1B, C). This finding complements our earlier model derived from bulk phenotypic and functional analyses [43] and places premature hematopoietic aging in HO-1–deficient mice into a broader framework of stem cell exhaustion driven by imbalanced activation rather than uniform overproliferation. Notably, in aged HO-1 knockout mice, hematopoiesis appears to be maintained by a minor subset of active stem and progenitor cells, while the majority of HSCs remain deeply quiescent (Fig. 1B).

This revised view also explains the seemingly paradoxical observation that aged HO-1– deficient HSCs exhibit lower levels of DNA damage than their wild-type counterparts. Our previous comet assay analyses show that while young HO-1–deficient HSCs accumulate elevated DNA breaks [43], aged HO-1–deficient HSCs display reduced DNA damage relative to age-matched controls (Fig. 2A). We propose that this reflects a shift toward a largely non-cycling stem cell population in aged knockout mice, which may be protected from replication-associated DNA lesions despite an intrinsically compromised DNA damage response [56]. In contrast, young HO-1–deficient HSCs exhibit increased susceptibility to genotoxic stress, as further supported by the altered mutational landscape observed after low-dose irradiation (Fig. 2C).

Mechanistically, we show that HO-1 deficiency disrupts multiple components of the G1/S checkpoint in HSCs, including impaired activation of Chk2, altered regulation of Cdc25A and Cdk1 phosphorylation, and increased replication origin licensing marked by elevated Cdc45-high cells (Figs. 3D–F, 4A). These defects are accompanied by reduced nuclear Ku70 signal intensity in HSCs, despite elevated DNA damage (Fig. 4D, E). Given the central role of Ku70 in early DNA double-strand break sensing and checkpoint signaling [57–59], this defect likely contributes to the failure of HO-1–deficient HSCs to properly arrest the cell cycle or undergo apoptosis in response to genotoxic stress. Importantly, the observation that Ku70 and 53BP1 signals are increased in more differentiated progenitors suggests that HO-1 regulates DNA damage responses in a cell-type–specific manner along the hematopoietic hierarchy (Fig. 4 D, E). It is worth noting that in a previous affinity purification–mass spectrometry analysis performed in HEK293 cells, Ku70 was identified as a candidate component of nuclear HO-1–containing protein complexes under hypoxic conditions [36].

One of the most striking findings of our study is that HO-1 exhibits a predominantly nuclear localization in LT-HSCs under steady-state conditions. Confocal imaging and subcellular analyses demonstrate that nuclear HO-1 is abundant in LT-HSCs but progressively redistributed toward the cytoplasm upon early differentiation into multipotent progenitors, a process that becomes more pronounced with aging (Fig. 5A–C). These observations argue against nuclear HO-1 being merely a stress-induced artifact and instead support the idea that it fulfills a dedicated regulatory function in stem cells.

By employing an inducible *Hoxb5*-CreERT2 model, we were able to dissect the HSC-intrinsic role of HO-1 with unprecedented specificity (Fig. 6A). Importantly, deletion was induced in young mice and analyzed after physiological aging, allowing us to uncouple intrinsic HO-1 functions from the inflammatory and niche-mediated effects that dominate in global HO-1 deficiency [43]. Transcriptomic profiling of HO-1–deficient LT-HSCs revealed broad dysregulation of pathways involved in translation, mitochondrial metabolism, and differentiation (Fig. 6B–E; Supplementary Figs. 1–2). These changes were consistent with increased cellular activation and early lineage priming, and were functionally validated by enhanced clonogenic output and accelerated early differentiation in single-cell assays (Fig. 6F, G). Notably, despite these changes, lineage output was not overtly skewed, indicating that HO-1 primarily restrains the earliest commitment steps and controls the activation threshold rather than dictating terminal fate decisions. This pattern is consistent with functional heterogeneity within the HO-1–deficient HSC compartment. Importantly, these results complement our previous studies showing that myeloid bias of HSCs in HO-1^-/-^ mice is at least in part enhanced by HO-1 deficiency in the bone marrow niche [41], thereby delineating distinct intrinsic and extrinsic levels of HO-1–dependent regulation.

The transcriptional signatures observed in HSC-intrinsic HO-1 deletion differ substantially from those previously reported in HSCs isolated from global HO-1 knockout mice [43], where inflammatory signaling and niche-derived stress dominate. This distinction underscores the importance of context: while global HO-1 deficiency profoundly alters the hematopoietic microenvironment, our data reveal that even in an otherwise normal niche, intrinsic HO-1 activity is required to preserve HSC quiescence, genome integrity, and differentiation timing during aging.

Although our study establishes HO-1 as a nuclear regulator of HSC function, the precise molecular mechanisms underlying this activity remain to be defined. While we did not observe global enrichment of DNA damage response pathways at the transcriptomic level (Fig. 6D), reduced expression of Xrcc6 (Ku70) in HO-1–deficient LT-HSCs suggests a direct intrinsic effect on genome maintenance pathways (Fig. 6G). Whether nuclear HO-1 directly modulates DNA repair machinery, chromatin organization, or transcription factor activity remains an important question for future studies.

Importantly, in our previous work we demonstrated that HO-1 deficiency increases susceptibility to replication stress in hematopoietic stem and progenitor cells (HPSCs) isolated from murine bone marrow and cultured ex vivo [37]. Proliferating HO-1–deficient HSPCs exhibited a higher proportion of stalled replication forks, while the mean length of ongoing forks was increased. Across multiple cell types, we further observed that in the absence of HO-1 the response to DNA conformational hindrance less stringent. This was accompanied by accumulation of DNA G-quadruplex structures stabilized by excess heme and an impaired replication stress response regulated by the PARP1-p53-p21 axis [37]. These findings suggest that protection against replication stress may represent a general component of the cytoprotective activity of HO-1. Although not directly addressed in the present study, a similar mechanism may operate in HSCs. This possibility is indirectly supported by our previous observation of nuclear colocalization of HO-1 with DNA G-quadruplexes in HSCs isolated from mouse bone marrow, as detected by PLA [60].

Our study limitation is that while the *Hoxb5*-CreERT2 model provides one of the most specific genetic tool currently available to target functional LT-HSCs, *Hoxb5* expression and, therefore, the Cre expression as well, it is restricted to a subset of phenotypic LT-HSCs [61]. As a result, HO-1 deletion likely occurred in only a fraction of the LT-HSC population analyzed, potentially underestimating the full magnitude of the intrinsic phenotype. Nevertheless, the robust transcriptional and functional effects observed despite partial targeting highlight the sensitivity of early HSC fate decisions to HO-1 activity.

In summary, our findings redefine HO-1 as a nuclear, cell-intrinsic regulator of HSC homeostasis that restrains premature activation, preserves genome integrity, and supports proper differentiation dynamics during aging. By integrating proliferation assays, DNA damage analyses, subcellular localization studies, and precise genetic targeting, our study provides a comprehensive framework for understanding how non-canonical HO-1 functions shape stem cell behavior.

## Materials and methods

### Mice

All animal procedures were performed following national and European legislation and approved by the Second Local Ethics Committee on Animal Testing in Kraków (approvals number 113/2014, 47/2019, 120/2019, 121/2019, 342/2020, 83/2021). Experiments were performed on 3-4-month-old mice from the in-house colony of strain C57Bl/6×FVB Hmox-1 WT and KO (HO-1^-/-^ and HO-1^+/+^). The old group of mice used in some of the experiments were at least 1 year old. The Hmox-1^flox/flox^ mice were maintained on C57BL/6 background. The C57BL/6xFVB HO-1^-/+^ and Hmox-1^fl/fl^ mice were kindly provided by Dr. Anupam Agarwal, University of Alabama, Birmingham, USA. The *Hoxb5*-CreERT2 mice were kindly provided by Dr Irving Weissman, Stanford University, USA. The CreERT2 construct were introduced in the same loci as in the Hobx5-mCherry mice, as described previously [61], by Transgenic, Knockout, and Tumor Model Center at Stanford University.

To induce Cre recombinase in Hmox-1^flox/flox^;*Hoxb5*-CreERT2 mice were administered with tamoxifen (Sigma-Aldrich, dissolved in corn oil at concentration of 20 mg/ml) by intraperitoneal injections at 75 mg/kg body weight for 5 consecutive days.

### Flow cytometry analysis and cell sorting

Flow cytometry analysis was done on LSR Fortessa cytometer (BD Sciences). Cell sorting was done on a MoFlo XDP cell sorter (Beckman Coulter). Unless otherwise indicated, the populations used in studies were defined as follows: LT-HSCs – LSK CD150^+^CD48^-^CD34^-^, ST-HSCs – LSK CD150^+^CD48^-^CD34^+^, MPP – LSK CD150^-^CD48^+^. Following antibody clones were used in the study: Lin cocktail (clone 17A2/RB6-8C5/RA3-6B2/Ter-119/M1/70, BioLegend), Sca-1 (clone D7, ThermoFisher Scientific), c-Kit (clone 2B8, ThermoFisher Scientific), CD150 (clone TC15-12F12.2, BioLegend), CD48 (clone HM-48-1, BioLegend), CD34 (clone RAM34, BD Biosciences).

For intracellular staining, no more than 10 million cells were taken per sample, with an even cell count between samples, including controls. After counting, cells were stained with surface antibodies as usual and then, without a washing step, subjected to the IntraSure kit (BD Biosciences) procedure. In short, 200 µl of reagent A were added to each sample right after the staining, samples were vortexed and incubated for 5 min in darkness, RT. Next, 2.5 ml of BD FACS Lysing Solution diluted in sterile water was added to each sample. Samples were once again vortexed and incubated for another 10 min, RT, in darkness. After incubation time, samples were centrifuged (800 x g, 5 min) and the supernatant was discarded. Cells were resuspended in 100 µl of an antibody of interest against an intracellular target diluted in reagent B. Samples were incubated for 45 min, RT, in darkness. If the staining required a secondary antibody after washing cells with PBS w/o, the step with reagent B was repeated. Samples were washed in PBS w/o and resuspended in PBS w/o with DAPI (to distinguish nucleated cells from debris) for collection on the flow cytometer. Following antibody clones were used for the intracellular staining: Cdc25A (clone F-6, Santa Cruz Biotechnology), Cdc25A Ser124 (Bioss Antibodies), Cdc25A Ser76 (Biorbyt), Cdc45L (clone JJo91-04, ThermoFisher Scientific). Cdk1 (clone 17, Santa Cruz Biotechnology), Cdk1 Tyr15 (clone E.658.6, ThermoFisher Scientific), Cdk2 (clone D-12, Santa Cruz Biotechnology), Cdk2 Thr160 (Cell Signaling), Chk1 (clone G-4, Santa Cruz Biotechnology), Chk1 Ser345 (clone R3F9, Abnova), Chk2 (clone A-12, Santa Cruz Biotechnology), Chk2 Thr68 (clone ebchk2, eBioscience), Mdm2 (clone D-7, Santa Cruz Biotechnology), Mdm2 Ser166 (Biorbyt).

### Annexin V assay

Annexin V assay was performed using a TACS Annexin V kit from Trevigen. In short, cells were counted on Muse Cell Analyzer after isolation and lysis of red blood cells. Five million cells were taken for each sample, stained with surface markers, and washed. Pelleted cells were resuspended in 100 µl of Annexin V Incubation Reagent, which constituted of 10 µl 10X Binding Buffer, 1 µl Annexin V-FITC, and 89 µl dH2O per sample. One mix was prepared for all the samples. Samples were incubated in the dark for 15 min, RT, and then 400 µl of 1X Binding Buffer was added. Samples were processed on a flow cytometer within 1 hour of staining.

### BrdU assay

HO-1^+/+^ and HO-1^-/-^ adult mice (3-month-old) were injected peritoneally with 100 mg BrdU (Sigma Aldrich) per kg body weight in PBS w/o as a starting time-point. Animals were maintained on 1 mg/ml BrdU in the drinking water with free access throughout the study. Water bottles were protected from light and changed every 7 days. Mice were sacrificed at different time points for subsequent flow cytometry analysis.

### Alkaline Comet Assay

Alkaline comet assay was performed with the Trevigen CometAssay kit according to the manufacturer’s protocol. In brief, LT-HSCs from either young (10–12-week-old) or aged (>18-month-old) HO-1^-/-^ and HO-1^+/+^ mice were sorted into a small volume of PBS w/o, embedded in Comet LMAgarose, and transferred onto a single CometAssay HT Slide. The slide was then incubated at 4°C for 30 min to enhance the gelling process and placed in Lysis Solution for overnight incubation at 4°C. Next, cells were treated with a freshly made alkaline unwinding solution for 20 min, RT, and subjected to electrophoresis in alkaline conditions with an adjustment for electrophoresis units other than supplied by Trevigen (4°C, 1 Volt/cm, 300 mA, 30 min). After electrophoresis, the slide was washed twice in dH2O followed by a single 70% ethanol wash, and dried at 37°C. Samples were stained with SYBR Gold and imaged. Analysis was performed on blinded files with either CometScore (TriTek) or CaspLab software, which calculated values for the percent of Tail DNA and Olive Tail Moment.

### Analysis of the HO-1 localization on sorted cells

The analysis of HO-1 protein localization was done by sorting the cells on poly-L-lysine coated microscopic slides in custom-defined areas. Next, cells were let to settle down and fixed with 2% PFA, and stained according to immunocytochemistry protocol with SPA-896 antibody (ENZO, diluted 1:200) and Alexa488 goat anti-rabbit (Invitrogen, diluted 1:400). Cells were image with LSM710 confocal microscope with 63x immersion objective (Carl Zeiss). The nuclear and cytoplasmic signal was analyzed by Image J, by hand-made masks for the whole cell area based on brightfield image and nucleus based on DAPI signal. The masks were then used to analyze the HO-1 signal intensity.

### RNA-seq analysis

Each sample represents HSCs sorted and pooled from two mice. 1000-3000 cells per sample were sorted directly into lysing buffer from Single Cell RNA Purification Kit (Norgen, Biotek), and RNA isolation was performed according to the manufacturer’s protocol. The isolation of the required RNA amount and quality was confirmed by Bioanalyzer PicoRNA Kit (Agilent).

The library was performed according to the SmartSeq2 protocol described by Picelli et al. [62,63]. In short, the strategy relies on template switching based on LNA primers that bind to the end of the 3’ cDNA transcript. cDNA is amplified by PCR (12-14 cycles were required depending on the amount of the RNA) and purified by Ampure XP beads (Beckman Coulter) with a 1:0.6 (DNA/beads) ratio. The amount of obtained amplified transcriptome was quantified using Bioanalyzer with High Sensitivity DNA Chip (Agilent) and 200 pg of input material was used to Nextera XT Kit (Illumina) for tagmentation, adapter ligation, and final PCR enrichment. The obtained libraries were quantified by High Sensitivity DNA Chip and sequenced by 75 bp pair-end reads using NextSeq500 (Illumina). The reads were pseudo-mapped to mm10 reference with Salmon or Star software [64,65]. The number of mapped reads ranged from 30.4 to 53.8 million reads per sample.

The analysis of expression of a particular gene and statistical analysis was done using the DESeq2 package in R [66]. The GeneOntology analysis was done using the ClusterProfiler package [67].

### Whole-exome sequencing (WES)

HO-1^+/+^ and HO-1^-/-^ adult mice (3-month-old) were divided into two groups each: control group and a group that received a low dose irradiation (2 Gy, Cs-137 source). Mice were left to age for a period of 6 months and their bone marrow was harvested and DNA was isolated by standard column-based protocol. The WES was performed using Twist Bioscience panel by EMBL Genomics Core Facility, Heidelberg, Germany.

Pair-end sequences in fastq format for all 16 samples were aligned to GRCm39 reference genome using the Dragen open-source mapper/aligner (DRAGMAP v1.3.2). Alignment results were postprocessed by marking duplicates with samblaster v.0.1.26 [68] and sorted with sambamba v.1.0.1 [69]. The variant calling workflow was split between 4 germline samples (wild-type non-radiated) and 12 somatic samples (knock-out irradiated, knock-out non-radiated and wild-type irradiated). Variants for germline samples were called with HaplotypeCaller (GATK 4.6.0.0 [70] with default parameters and enabled dragen-mode. Variants detected in germline samples were used for creating control panel (“normals”) to capture somatic alterations in our 12 somatic samples. Both creating of panel of normals and variant calling for somatic samples was conducted using Mutect2 (GATK 4.6.0.0 [70]). Consequences of all variants called in germline and somatic workflow were annotated with Ensembl VEP 113 [44] and their impact was classified based on their severity as LOW, MODERATE or HIGH.

To determine whether genotype affected the distribution of mutation-impact severity after irradiation, variants were annotated and grouped into LOW-, MODERATE-, and HIGH-impact categories. For statistical comparison of mutation-impact distributions between conditions, 2×3 contingency tables (Group × Impact category) were constructed. Overall distribution differences were evaluated using a chi-square test of independence. To identify which mutation classes drove the shifts, individual 2×2 Fisher’s exact tests were performed for each impact category by comparing the number of mutations in that class versus all other classes combined. Results with p < 0.05 were considered significant.

### Real-time PCR analysis

Reverse transcription (RT) was performed using 1 μg of RNA and the RevertAid Reverse Transcriptase kit (ThermoFisher). In brief, 1 μl of oligo(d)T primers (100 μM) was added to the RNA sample and incubated for 10 minutes at 72 °C. Next, 2.9 μl of a reaction mixture containing dNTPs (10 mM), 5× RT buffer, and RevertAid reverse transcriptase (200 U/μl) was added. The samples were then incubated for 1 hour at 42 °C.

Quantitative real-time PCR (qRT-PCR) was used to assess the expression levels of the studied genes. cDNA obtained in the previous step was diluted fivefold, and 3 μl of each sample was transferred to a MicroAmp™ Optical 96-Well Reaction Plate (ThermoFisher). Each reaction contained 7.5 μl JumpStart SYBR Green PCR Master Mix (Sigma-Aldrich), 0.75 μl of forward primer (10 μM), 0.75 μl of reverse primer (10 μM), and 3 μl of water. The sequences of the primers are listed in table below (Table 1). The gene expression was assessed on a StepOnePlus thermocycler (Applied Biosystems, Waltham, MA, USA).

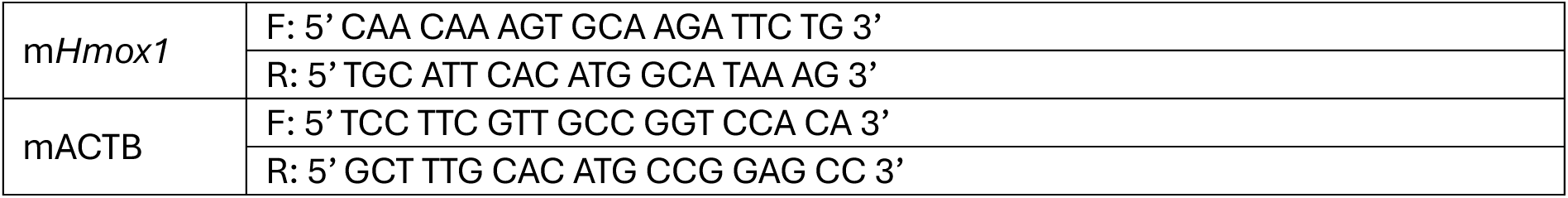

### Statistical analysis

Statistical significance for all assays was calculated using GraphPad Prism software. All results were analyzed for the normal distribution with D’Agostino-Pearson omnibus and Shapiro-Wilk normality tests. Statistical significance between 2 groups was determined using a paired or unpaired Student’s t-test, as appropriate. Mann-Whitney test was used for the data with non-normal distribution. When the experimental scheme included two variables e.g. time and genotype, a two-way ANOVA test was performed. In this case, the statistical significance of a given variable was shown only when no significant interaction between variables was detected. Results with p < 0.05 were considered significant. * p = 0.01 to 0.05, ** p = 0.001 to 0.01, *** p = 0.0001 to 0.001, **** p < 0.0001.

## Acknowledgments

We would like to acknowledge Agnieszka Andrychowicz-Rog and Joanna Uchto for technical support. The study was supported by the National Sciene Center (grant Harmonia no NCN2015/18/M/NZ3/00387 awarded to A.J., grant Preludium no NCN 2017/25/N/NZ1/02156 and ETIUDA 2019/32/T/NZ3/00624 awarded to M.Z.), Foundation for Polish Science (Fellowship START 100.2020 awarded to M.Z) and European Research Council (Starting Grant “StemMemo” nr 101041737 awarded to KS).

The research has been supported by grants from the Priority Research Area BioS and the Faculty of Biochemistry, Biophysics and Biotechnology (FBBB) under the Strategic Programme Excellence Initiative at Jagiellonian University (JU).

Presented results were a part of M.Z.’s PhD thesis.

During the preparation of this work, the authors used ChatGPT in order to check and correct language and readability. The authors reviewed and edited the content as needed and take full responsibility for the content of the publication.

## Supplementary Figures 1-3

Representations of significantly enriched Gene Ontology (GO) biological processes in gene set enrichment analysis (GSEA) in LT-HSCs isolated from HO-1^ΔHSC^

---

**Supplementary Figure 1.**
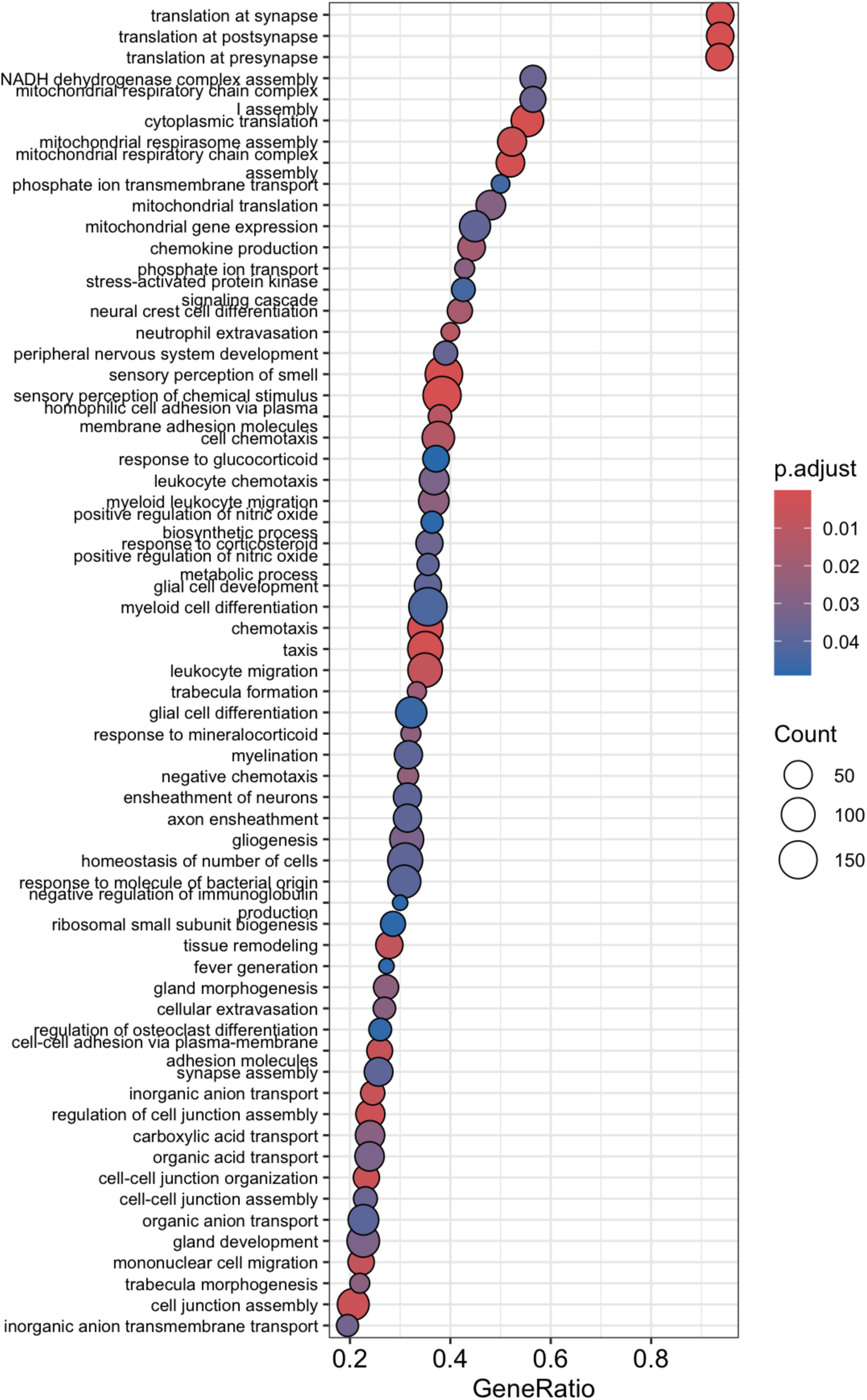

**Supplementary Figure 1.**
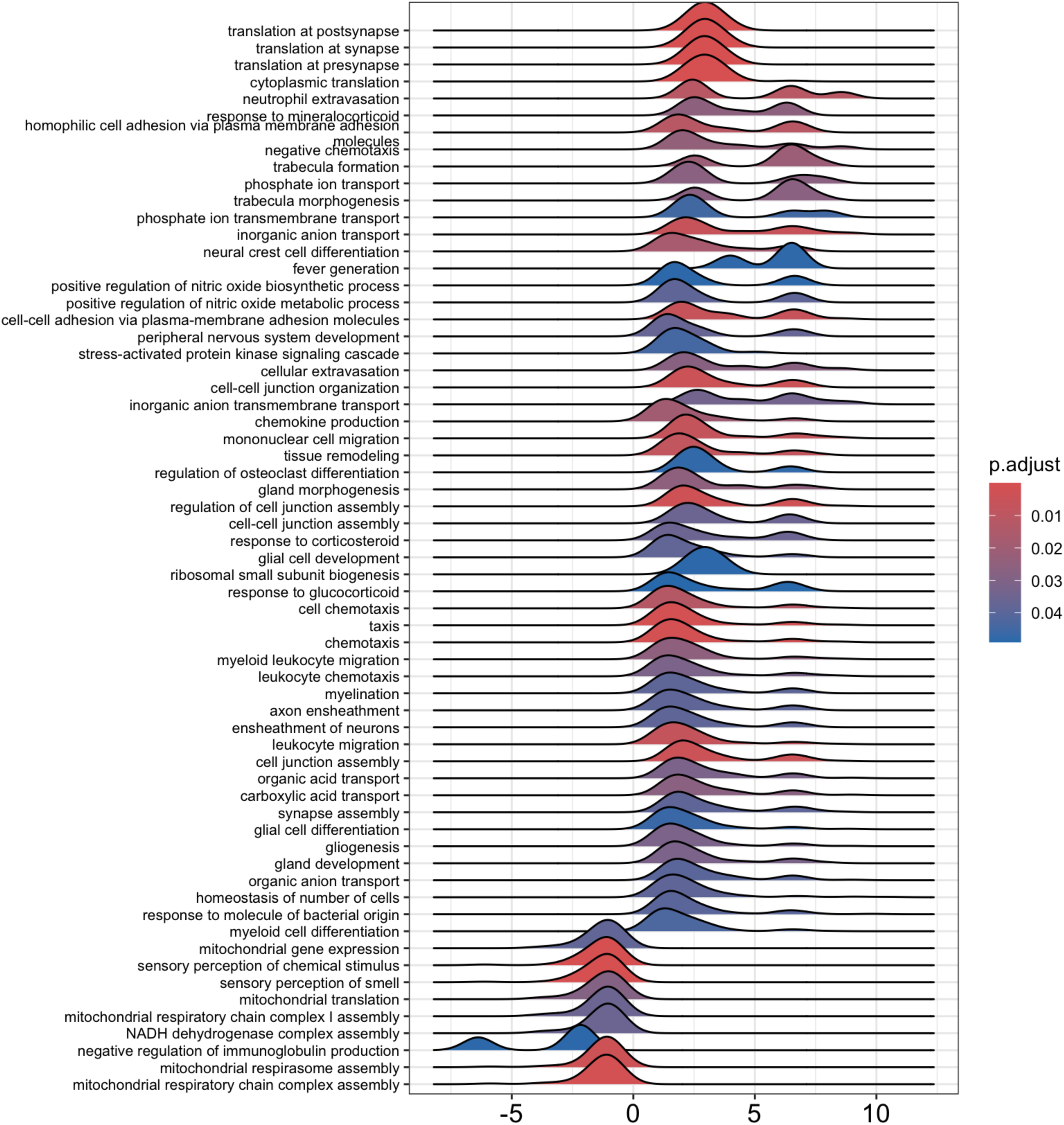

**Supplementary Figure 1.**
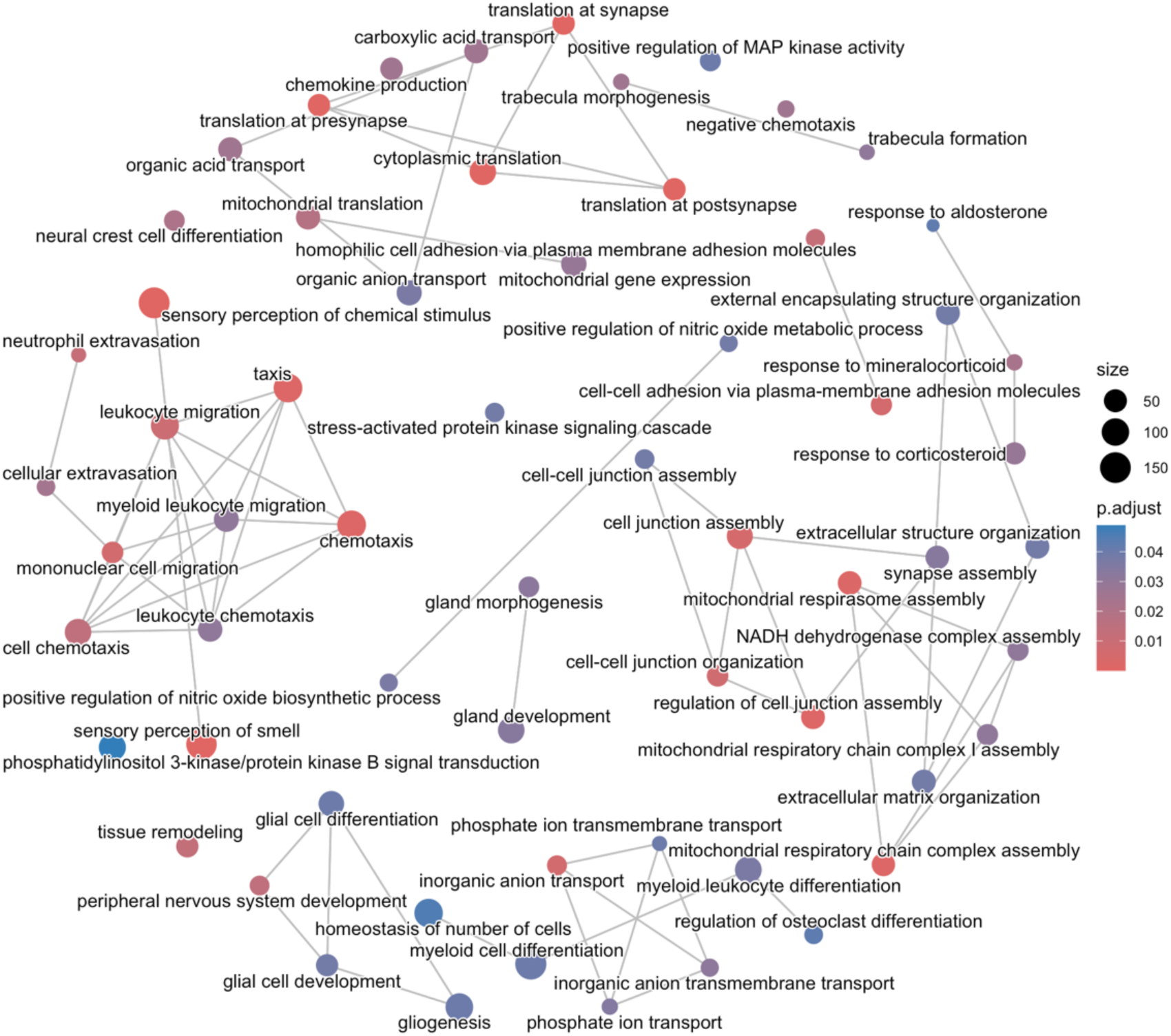

**Supplementary Table 1.**

| IMPACT | CONSEQUENCE | HO-1<sup>+/+</sup><br>irradiated | HO-1<sup>-/-</sup> non-<br>irradiated | HO-1<sup>-/-</sup><br>irradiated |
| --- | --- | --- | --- | --- |
| LOW | splice donor 5th base variant | 48 | 48 | 55 |
| LOW | splice donor region variant | 178 | 160 | 204 |
| LOW | splice polypyrimidine tract<br>variant | 708 | 692 | 831 |
| LOW | splice region variant | 899 | 819 | 930 |
| LOW | stop retained variant | 32 | 20 | 24 |
| LOW | synonymous variant | 5466 | 5188 | 6175 |
| MODERATE | inframe deletion | 114 | 88 | 111 |
| MODERATE | inframe insertion | 55 | 61 | 61 |
| MODERATE | missense variant | 7221 | 6747 | 7303 |
| MODERATE | protein altering variant | 3 | 2 | 5 |
| HIGH | frameshift variant | 272 | 242 | 248 |
| HIGH | splice acceptor variant | 72 | 83 | 59 |
| HIGH | splice donor variant | 91 | 79 | 97 |
| HIGH | start lost | 112 | 114 | 111 |
| HIGH | stop gained | 351 | 341 | 337 |
| HIGH | stop lost | 76 | 75 | 71 |

